# Modeling fragment counts improves single-cell ATAC-seq analysis

**DOI:** 10.1101/2022.05.04.490536

**Authors:** Laura D. Martens, David S. Fischer, Vicente A. Yépez, Fabian J. Theis, Julien Gagneur

## Abstract

Single-cell ATAC-sequencing (scATAC-seq) coverage in regulatory regions is typically binarized as an indicator of open chromatin. However, the implications of scATAC-seq data binarization have not systematically been assessed. Here, we show that the goodness-of-fit of existing models and their applications, including clustering, cell type identification, and batch integration, are improved by a quantitative treatment of the fragment counts. These results have immediate implications for scATAC-seq analysis.

---

Single-cell ATAC-seq (scATAC-seq^1^) is a major method employed to study chromatin regulation in health and diseases. scATAC-seq leverages the fact that most active cis-regulatory regions are accessible^2^. In the ATAC-seq (assay for transposase-accessible chromatin using sequencing) protocol, the prokaryotic Tn5 transposase is used to insert sequencing adapters into accessible regions of the genome^1,3^. Reads in scATAC-seq data reflect Tn5 insertion events in individual cells^1^ (Fig. 1a, example Fig. 1b). When analyzing scATAC-seq data, open chromatin regions are generally first called on the pooled data as peaks, i.e., genomic regions with a significant excess of reads compared to background^1,4,5^. Alternative approaches define the feature set as genomic windows or bins^5,6^. Once the feature set is defined, the reads overlapping each feature are counted for each cell. The resulting count matrix is typically very sparse, with less than 10% non-zero counts^7^.

**Figure 1.**
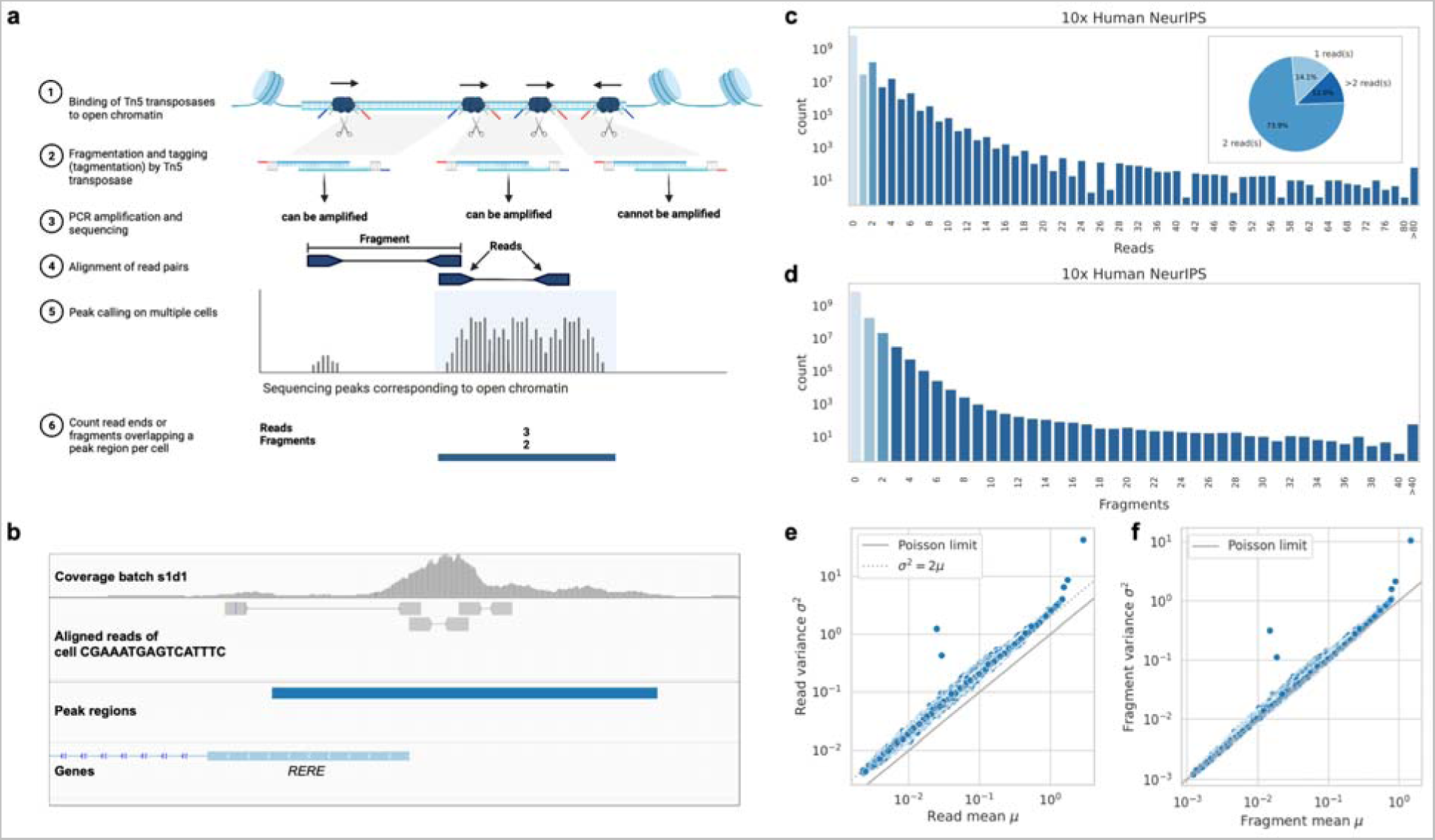
scATAC-seq data is quantitative. **a)** Illustrated is the scATAC-seq protocol and count aggregation strategy. Tn5 transposases insert into open chromatin regions, cut the DNA, and attach sequencing adapters (blue and red). Two Tn5 insertions create one fragment with adapters. The orientation of insertion is important as only fragments flanked with two distinct barcodes can be captured and amplified. Fragments are sequenced paired-end and aligned to the genome. scATAC-seq peak calling is performed using reads from multiple cells. Once peak regions are identified, reads (deduplicated fragment ends) or fragments overlapping the peak region are counted for each cell separately. **b)** Genome viewer snapshot of one peak region in the NeurIPS dataset at the promoter of the human gene *RERE* showing multiple insertions in a single cell. The tracks show, from top to bottom, the coverage of one batch used for peak calling, the aligned read pairs of a single cell, the peak region, and genome annotation. The peak region overlaps with five reads and three fragments. **c)** Read count distribution on the entire NeurIPS dataset. The striking odd/even pattern in read count distribution reflects that reads come in pairs and suggests that fragment counts, rather than reads, should be modeled. Inset: A pie chart showing the percentage of all non-zero peaks with 1, 2, or more than 2 reads. **d)** Distribution of the approximated fragment count does not show an even/odd pattern. **e)** Variance of read counts across cells against mean read counts. Each dot represents one peak region. When fragment ends (reads) are counted, the variance of read counts is about twice the mean (grey dotted line), which is not consistent with a Poisson distribution (solid grey line). **f)** Same as (e), but for fragment counts. The variance of fragment counts is approximately equal to the fragment count mean, consistent with a Poisson distribution (solid grey line).

Machine learning modeling of scATAC-seq data is used to support biological investigations of single-cell genome regulation, including identification of cell types, differentially accessible regions, and inference of transcription factor activity, among others. Irrespective of the modeling strategy and the application purpose, one guiding criterion for fitting any machine learning model is the choice of a loss function that captures the goodness-of-fit of a model to the data. Therefore, the data representation and the choice of the loss function are of prime importance for all those models. Due to overall data sparsity and a conceptualization of chromatin accessibility as a binary state, it is customary in many methods to binarize the read count matrix by default prior to machine learning modeling of scATAC-seq data^5–10^. While handling the data quantitatively has been done in some methods^4,11,12^, there is no systematic investigation of the advantages or disadvantages of binarizing the data.

Here we study the effects of binarization versus count-based models on scATAC-seq data modeling tasks and assess the quality of the learned latent space using multiple downstream methods. We based our analysis on four publicly available datasets representing different protocols, species, and tissues, each with cell-type annotation and several batches (Supplementary Table 1, Online Methods): (i) a 10x multiome single-cell RNA-seq and ATAC-seq human bone marrow dataset^13^ (“NeurIPS”, 62,501 cells) (ii) a 10x human hematopoiesis dataset consisting of flow-sorted and unsorted samples from bone marrow and blood^14^ (“Satpathy”, 63,882 cells) (iii) a 10x scATAC-seq profile of the entire fly brain^15^ (“Fly”, 117,613 cells) (iv) a large sci-ATAC-seq3 dataset of 53 fetal tissue samples representing 15 organs^16^ (“sci-ATAC-seq3”, 720,613 cells).

We first considered the proportion of peaks above the typical binarization threshold of one read. Across all datasets, more than 65% of non-zero peaks had more than one read count (Fig. 1c and Extended Data Fig. 1). For example, in the NeurIPS dataset, as many as 74% of non-zero peaks had counts of two and 12% had even more than two reads. When studying the distribution of counts in more detail, we found that there were 5-fold more peaks with even than odd counts. This pattern can be explained as an artifact of the count aggregation strategy: In the 10x Genomics Cell Ranger ATAC pipeline^17^, reads (deduplicated fragment ends) are counted instead of the fragments themselves because they mark transposase insertion events (Fig. 1a). Since scATAC-seq creates paired-end reads, this will mostly result in even counts, whereas odd counts are only generated if one read pair lies outside the peak region (Fig. 1a,b). The read count matrices from the 10x pipeline are the basis for many downstream processing tools^4,7–10^. Another widely used tool, ArchR^5^, also counts read ends and binarizes them by default. On the other hand, the inbuilt counting function in Signac^4^ counts fragments. However, Signac can also work directly with the 10x read count matrix. Despite the different approaches and inconsistencies employed in scATAC-seq analysis methods, there has not been a benchmark evaluating the read versus fragment count strategy.

In machine learning, loss functions are typically based on the choice of a parametric distribution of the data. However, the alternating pattern of odd and even read counts is not trivially represented by common statistical distributions. In contrast, fragment counts - estimated here from the existing pipelines by simply aggregating odd and neighboring even reads - showed a regular monotonic decay (Fig. 1d and Extended Data Fig. 1; Online Methods). Count observations are commonly modeled using the Poisson distribution, for which the mean equals the variance. We found that the variance of read counts for each region across cells was approximately twice the mean (Fig. 1e and Extended Data Fig. 1), violating the Poisson distribution assumption and consistent with the fact that reads typically come by two. In contrast, the mean-variance relationship of fragment counts was broadly consistent with a Poisson distribution across the four datasets and the entire dynamic range (Fig. 1f and Extended Data Fig. 1).

Altogether, these results have two implications. First, scATAC-data contains information beyond a binary accessibility state. Second, fragment counts, but not read counts, can be more suitably modeled with the Poisson distribution.

Next, we investigated the effects of modeling the fragment counts rather than a binarized signal on downstream applications. We first investigated latent space learning of scATAC-seq data, a key first step for cell type identification, batch integration, and trajectory learning. To this end, we adapted the PeakVI model, a state-of-the-art variational autoencoder for scATAC-data^9^, to model the fragment count matrix. PeakVI originally models binarized ATAC data by learning a Bernoulli probability that a peak in a given cell is accessible. PeakVI corrects for cell-specific effects and region biases by learning cell-specific and region-specific factors. We modified PeakVI to model Poisson-distributed data in a model that we call Poisson VAE (Poisson variational autoencoder). To directly compare the choice of loss function, we used the same architecture as PeakVI up to the last layer, where we modeled a Poisson rate parameter for a given peak region in a cell (Online Methods). The total number of fragments per cell varies drastically across cells (Extended Data Fig. 2a). We included total fragment count as a pre-computed offset in the loss instead of learning a cell-specific factor, as done in PeakVI. Similarly, we tested the effect of using the precomputed offset in the binary case (Binary VAE) by including the proportion of non-zeroes in the binary loss function (Online Methods).

We first assessed how well the models fitted the four studies. As a common ground, we benchmarked the models for predicting the presence of at least one read, which is the typical binarization threshold. As scores, we used for the binary models the predicted probability of a region being open. For the quantitative models, we converted the prediction into a probability of having a count greater than one (Online Methods). Poisson VAE significantly outperformed PeakVI in reconstructing binarized counts as measured by average precision on three of the four datasets (NeurIPS: P = 1.2 × 10^−7^, Satpathy: P = 6.9 × 10^−8^, Fly: P = 3.7 × 10^−6^, Fig. 2a). Most of the performance boost could be attributed to using the observed total fragment counts rather than predicting it since the binary model (Binary VAE) also showed significantly better reconstruction than PeakVI. The quantitative model Poisson VAE significantly outperformed the binary model Binary VAE (NeurIPS: P = 1.3× 10^−6^, Satpathy: P = 7.1× 10^−3^, Fly: P = 7.1 × 10^−6^), suggesting that there is useful information in the quantitative data. We further tested that the performance improvement was not a result of disproportionately giving more weight to regions with high counts (Extended Data Fig. 2b). In the sci-ATAC-seq3 dataset, we did not observe a benefit from using quantitative counts or the observed total fragment count. The sci-ATAC-seq3 dataset (based on combinatorial indexing) has, on average, only half of the fragments per cell and peak detected compared to the 10x datasets (median fragment count per peak 0.036 vs. 0.017, Extended Data Fig. 2a and Supplementary Table 1). Downsampling of the NeurIPS dataset confirmed that the benefits of the quantitative over the binary model increased with a higher total fragment count (Extended Data Fig. 2c). However, it does only partially explain the lower performance on the sci-ATAC-seq3 dataset so other factors like the increased coverage variability in that dataset (Extended Data Fig. 2a) could play a role.

**Figure 2.**
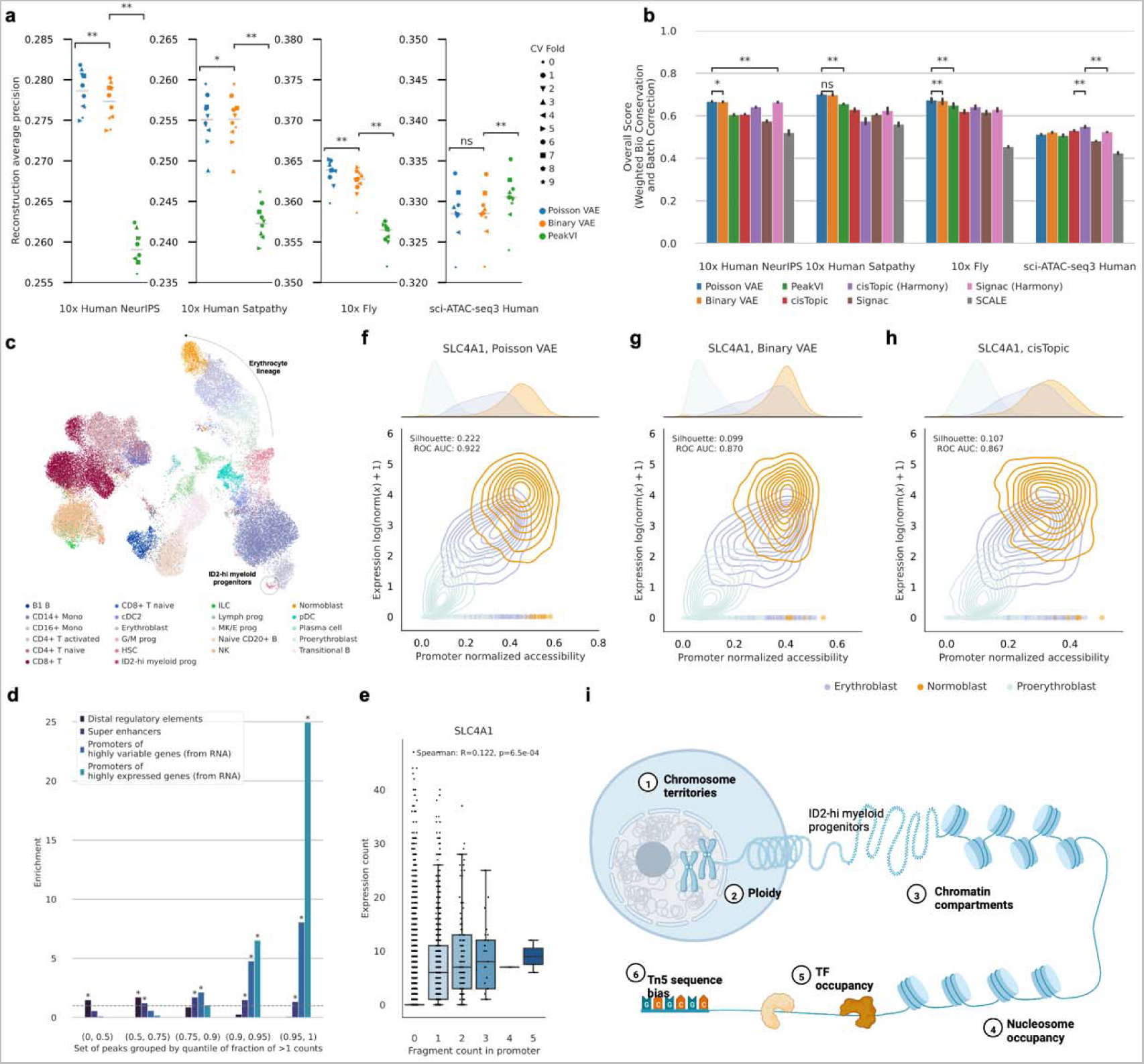
Modeling fragment counts instead of binarized scATAC data improves reconstruction, prediction, and latent representation of cellular profiles. **a)** Comparison of the Poisson VAE, Binary VAE, and PeakVI model on reconstructing the binarized cell-peak matrix of the NeurIPS, the Satpathy, the fly, and the sci-ATAC-seq3 datasets for ten cross-validation runs. Poisson VAE and Binary VAE use the observed total fragment count. Poisson rates were transformed to Bernoulli probabilities of generating more than one fragment, and performance wa evaluated using average precision. The horizontal line denotes the median. **b)** Comparison of integration accuracy for embeddings generated with Poisson VAE, Binary VAE, PeakVI, Signac, cisTopic, and SCALE on the datasets from (a). For cisTopic and Signac, additional batch correction was performed using Harmony. Overall integration accuracy scores were computed using a 40:60-weighted mean of batch correction and bio-conservation scores. **c)** UMAP of the integrated latent space of all NeurIPS batches, colored by cell type for the Poisson VAE model. The isolated label ID2-hi myeloid progenitors and the erythrocyte lineage are annotated. UMAPs for all other methods and datasets ar in Extended Data Fig. 5-8. **d)** Enrichment (odds ratio) of distal regulatory elements, super-enhancers in bone marrow, promoters of highly expressed genes and promoters of highly variable genes in the scATAC-seq peaks of the NeurIPS dataset. Peaks are sorted by the fraction of counts above the binarization threshold and grouped accordin to different quantiles. **e)** Correlation of expression of the *SLC4A1* gene and fragment counts in its promoter. The Spearman correlation is computed on cells with at least one fragment count in the promoter. We restricted the plot to cells of similar total fragment count (0.25-0.75 quantile) to not capture effects that total fragment count drives. We see a quantitative signal in promoter accessibility that would be lost by binarization. **f, g, h)** Log-normalized gene expression over normalized accessibility of the *SLC4A1* gene for the Poisson VAE (f), Binary VAE model (g) and cisTopic model (h). Cell type separation is measured with the silhouette width and area under the ROC curve and is better with the Poisson VAE model. **i)** Multiple biological factors contribute to DNA accessibility in single cells to be quantitative rather than binary. They include a diploid genome, density of chromatin packaging, nucleosome spacing, TFs in a peak region preventing the Tn5 from binding, and sequence preferences of Tn5. In all boxplots, the central line denotes the median, boxes represent the interquartile range (IQR), and whiskers show the distribution except for outliers. Outliers are all points outside 1.5 times the IQR. *P* values were computed using the paired Wilcoxon test. *P_□_<_□_0.05, **P_□_<_□_0.01, ***P_□_<_□_0.001, ****P_□_<_□_0.0001; Benjamini–Hochberg corrected. Error bars represent the 95% confidence interval over ten cross-validation runs.

We next investigated the implications of these improved fits for downstream applications, starting with batch integration. An ideal integration should reduce technical artifacts between batches while retaining biological variation. We evaluated several integration metrics divided into two categories, batch integration, and bio conservation, as previously described^18^. Four metrics fall into the first category and measure the mixture of batches in the resulting embedding. The second category has six metrics that measure the consistency of cell type annotation before and after integration, cell type separation, and isolated label preservation in the joint embedding (Online Methods). We used as ground truth proxy the provided cell type annotations. All four selected datasets were annotated using distinct approaches allowing for a more unbiased evaluation of model performance. Finally, an overall score is computed by weighting batch correction and bio conservation 40:60. In addition to the three VAE models, we compared embedding techniques of three other widely used methods. Signac^4^ and ArchR^5^ use latent semantic indexing (LSI) for dimensionality reduction. Since ArchR requires raw data as input, we used the Signac implementation, which takes read counts as input. cisTopic^8^ and SCALE^10^ work on the binarized read count matrix. cisTopic uses latent Dirichlet allocation (LDA) to learn a cell-topic matrix. SCALE is a deep generative model that implements a Gaussian Mixture Model in the latent space. Both Signac and cisTopic have an additional option to correct for batch effects using Harmony^19^.

Poisson VAE demonstrated the highest overall score in three datasets, with two of them showing a significant performance advantage over the second-ranked method (Fig. 2b and Extended Data Fig. 3). Moreover, these results were consistent across different weightings of bio conservation and batch correction metrics (Extended Data Fig. 4), suggesting that the improvements were not limited to batch correction only. For example, Poisson VAE better recovers the rare cell type ID2-hi myeloid progenitors in the NeurIPS dataset (only present in 2 of the 13 batches, Supplementary Table 1) as measured by the improved isolated label F1 score (Fig. 2c, Extended Data Fig. 3 and 5). Moreover, Poisson VAE and Binary VAE, which explicitly correct for the batch variable in the latent space, score highest for batch correction. For example, they successfully integrate the Kenyon cell subtype (KC-g) in the Fly dataset, in contrast to cisTopic and Signac (Extended Data Fig. 7). Harmony appears to overcorrect batch effects as it often leads to lower bio conservation scores (Extended Data Fig. 3 and 5-8). For the non-VAE models, cisTopic was consistently among the top four methods with good performance at bio conservation metrics.

We next explored what biological signal is in the quantitative data that could be captured in the Poisson VAE. We first investigated the characteristics of peaks that have a high number of cells with counts above the binarization threshold (Online Methods). We saw that peaks that fall in the top 5% based on the fraction of counts > 1 (high-count peaks) tend to be broader peaks (Extended Data Fig. 9). We further found that high-count peaks are enriched for promoter regions of highly expressed and highly variable genes across all cells of the dataset (as defined from matched scRNA-seq data) and super-enhancers, while low-count peaks are enriched for distal enhancer elements (Fig. 2d, Online Methods). This is in agreement with previous observations in bulk that active transcription start sites (TSS) are highly accessible and lie in regions of low nucleosome density while enhancers have lower accessibility^2^ and suggests that scATAC-seq also captures this accessibility continuum. We then investigated if increased accessibility at TSS also correlates with increased expression of the associated genes using the NeurIPS dataset, for which we had measurements of gene expression and accessibility in the same cell. To focus on the quantitative effect, we restricted the correlation computation to cells with at least one fragment count in the promoter region. We observed a significant correlation (Spearman corr., *P* < 0.05) of promoter accessibility with gene expression in 481 out of 3,879 genes (12.4%, 2.5 times higher than expected, Binomial Test *P* < 0.05). To investigate examples of this relationship, we considered cell type markers among the twenty genes with the highest correlation between promoter accessibility and gene expression (Extended Data Fig. 10a). One such example is *SLC4A1*, a gene involved in the red blood cell lineage^20^ (Spearman correlation: 0.12, *P*=0.001, Fig. 2c,e). Similarly, we found a significant correlation for genes involved in other biologically meaningful lineages (Extended Data Fig. 10b-d). We expected the Poisson VAE model to capture the quantitative accessibility signal and investigated if it was better at distinguishing cell types on these promoter regions. The normalized accessibility of Poisson VAE captured the gene expression trends of the cell types and showed improved cell type separation compared to CisTopic and Binary VAE as measured by the silhouette width and area under the ROC curve in three out of four cases (Fig. 2f-h and Extended Fig. 11, Online Methods).

In conclusion, we found that scATAC-seq data can be treated quantitatively and that useful information is lost through binarization of the counts. Modeling DNA accessibility in single nuclei quantitatively, rather than as a binary state, is consistent with the fact that chromatin accessibility is a highly dynamic and continuous state. Nucleosome turnover rates are in fact in the order of magnitude of the scATAC-seq incubation duration^1,21^. Moreover, transcription factors, just like transposase, have to diffuse through the nucleus to access DNA, likely reaching distinct chromosome territories and compartments with various efficiencies (Fig. 2h). Also, a single genomic position in diploid cells is not necessarily open or closed on both alleles synchronously. We saw that scATAC-seq fragment counts capture this continuum of chromatin accessibility. We observed that active promoter regions that are known to have high accessibility from bulk measurements^2^ also show high fragment counts in scATAC-seq. In parallel to our work and in agreement with our findings, a recent preprint also underscored the quantitative nature of scATAC-seq data and reported that gene expression increases with the number of fragments in accessible regions in single nuclei^22^. Moreover, we showed that fragment counts, but not read counts, can be modeled with the Poisson distribution. We found the scATAC-seq total fragment counts to be important for the reconstruction from scATAC data. However, we also observed that for the sparse sci-ATAC-seq3 dataset, there was less advantage of using the count data. This suggests that the quantitative treatment of scATAC-seq data could become even more relevant if the datasets become less sparse. Altogether, treating scATAC-seq data quantitatively is more general than binarization and it matters to study highly expressed and highly variable genes, which include some of the most important marker genes. These findings have immediate practical implications for modeling scATAC-seq data since using a Poisson loss over a binary loss comes at no noticeable increase in algorithmic complexity and computing time. Applications of this model to other settings such as multiome integration or other modalities that are typically binarized such as scChIP-seq^23^ could be the subject of future work.

## Methods

### Input data and preprocessing

#### NeurIPS Dataset

The multiome hematopoiesis dataset from the NeurIPS 2021 challenge^24^ was downloaded from the AWS bucket *s3://openproblems-bio/public/*. We did not perform any additional filtering of the data. scATAC-seq BAM files were downloaded from GEO (Accession GSE194122).

#### Satpathy dataset

The second hematopoiesis dataset^14^ was downloaded from GEO (Accession GSE129785). Specifically, the processed count matrix and metadata files: scATAC-Hematopoiesis-All.cell-barcodes.txt.gz, scATAC-Hematopoiesis-All.mtx.gz, scATAC-Hematopoiesis-All.peaks.txt.gz. We then filtered the peaks to only those that are detected in at least 1% of the cells in the sample, reducing the data from 571,400 to 134,104 peaks.

#### Fly dataset

Raw fragment files for chromatin accessibility of the fly brain^15^ were downloaded from GEO (Accession GSE163697). Additionally, peak regions, cell barcodes and cell metadata were extracted from the cisTopic object AllTimepoints_cisTopic.Rds which was downloaded from https://flybrain.aertslab.org. Fragments were counted per peak region using the Signac function FeatureMatrix. We then filtered the peaks to be detected in at least 1% of all cells. Furthermore, we excluded cells that were labeled unknown (CellType_lvl1 equal to ‘unk’ or ‘-’).

#### sci-ATAC-seq3 dataset

Count matrices and metadata were downloaded from GEO (Accession GSE149683)^16^. Peaks were filtered to be accessible in at least 1% of all cells.

### Fragment computation

The standard 10x protocol for generating the cell-peaks matrix is to count the fragment ends. As a simple way to estimate fragment counts, we rounded all uneven counts to the next highest even number and halved the resulting read counts.

### Poisson VAE model

Let *X*^*N×P*^ be a fragment count matrix consisting of *N* cells and *P* peak regions. We model the counts *x*_*cp*_with a variational autoencoder:

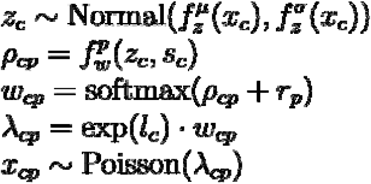

The encoder 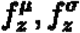 encode the parameters of a multivariate random variable *z*_*c*_ .*f*_*w*_ is a neural network that maps the latent representation *z*_*c*_ concatenated to the batch annotation *s*_*c*_ back to the dimension of peaks.*r*_*P*_ captures a region-specific bias such as the mean fragment count or peak length and is learned directly.*l*_*e*_ refers to the log-transformed total fragment counts per cell .*w*_*cp*_ is constrained to encode the mean distribution of reads over all peaks by using a softmax activation in the last layer. This means that.

### Binary VAE model

The binary VAE model models binarized counts:

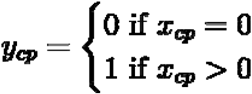

The binarized signal was modeled as follows:

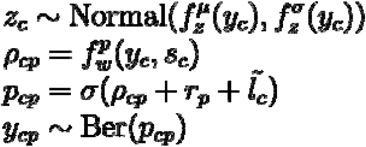

We included the proportion of non-zeroes by modeling:

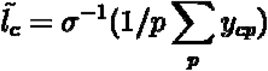

Here, σ^−1^ is the logit function. This way *P*_*cp*_ is equal to the mean accessibility of the cell for *P*_*cp*_ = *r*_*p*_ **=0**

### Encoder and decoder functions

The functions 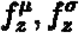 and the function *f*_*w*_ are encoder and decoder functions, respectively. To be as comparable as possible to PeakVI as implemented in scvi-tools^9,25^ (v.0.20.3), we use the same architecture. Specifically, these functions consist of two repeated blocks of fully connected neural networks with a fixed number of hidden dimensions set to the square root of the number of input dimensions, a dropout layer, a layer-norm layer, and leakyReLU activation. The last layer in the encoder maps to a defined number of latent dimensions *n*_*latent*_.

### Training procedure

We used the default PeakVI training procedure with a learning rate of 0.0001, weight decay of 0.001, and minibatch size of 128 and used early stopping on the validation reconstruction loss. We used a random training, validation, and test set of 80%, 10% and 10%, respectively. This was repeated ten times. We computed all evaluation metrics on the left-out test cells.

### Hyperparameter optimization

All models were run using the default PeakVI parameters. For the reconstruction task, we optimized the number of latent dimensions *n*_*latent*_ on the validation set for each dataset and model on reconstructing the binary accessibility matrix as measured by average precision. The used range was from 10 to 100 in increments of 10.

### Benchmarking methods

#### Signac

Count matrices were loaded into ChromatinAssays using Signac^4^ (v.1.9.0) and Seurat^26^ (v.4.3.0) without additional filtering (min.cells=min.features=0). We then computed the latent semantic indexing (LSI) embedding using the default procedure (RunTFIDF followed by RunSVD). We removed components that correlated with the total fragment count by more than 0.5. To investigate the effect of batch normalization, we created a batch-normalized LSI embedding by running RunHarmony with the respective batch variable as input.

#### cisTopic

We used the python implementation of cisTopic pycisTopic^8,27^ (v.1.0.3.dev2+g45b7e66.d20230426). cisTopic objects were created from the binarized count matrices. We then modeled the topics using the Mallet algorithm on 10 to 100 topics in steps of 10. We selected the optimal topic number using the suggested model selection metrics Minmo_2011^28^ and log-likelihood^29^. Finally, dimensionality reduction was performed on the cell-topic matrix with optionally first running Harmony to reduce batch effects.

#### SCALE

We used the provided Python script on https://github.com/jsxlei/SCALE to run SCALE^10^ on the binarized count matrix. We set the number of clusters to the number of cell types in the dataset. For visualization, a two-dimensional Uniform Manifold Approximation and Projection^30^ (UMAP) of the integrated latent space was generated based on the 15-nearest-neighbor graph. The cross-validation run with the best reconstruction was used.

### Evaluation

#### Reconstruction metrics

The reconstruction metrics were calculated on the binarized matrix. Poisson rate parameters λ_*cp*_ were transformed to a Bernoulli probability *P*_*cp*_ by computing the probability of getting one or more fragments in a peak for a given cell:

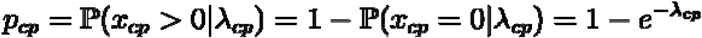

##### Average precision

As our reconstruction task is highly imbalanced (only a small fraction of all peaks are accessible), we used the average precision score as implemented in scikit-learn (v.1.2.2) to evaluate the reconstruction. Average precision estimates the area under the precision-recall curve.

#### Integration metrics

We used the scib^18^ (v.1.1.3) implementation for computing the integration metrics on the latent embedding of the cells. We used all available metrics using default parameters but excluded metrics that were specifically developed for scRNA-seq datasets (highly variable genes score, cell cycle score) and kBET due to its long run time. The trajectory score was only run for the NeurIPS dataset, which had a precomputed ATAC trajectory. Scib categorizes the metrics into metrics that measure batch correction and biology conservation.

Bio conservation comprises the following metrics that are applied to pre-defined cell-type labels that each dataset provided:

**Normalized mutual information (NMI):** measures the consistency of two clusterings. Here, we compare how well a clustering on the integrated embedding agrees with pre-defined cell-type labels. For optimal clustering, the scib package runs Louvain clustering at resolutions ranging from 0.1-2 in steps of 0.1. **Adjusted rand index (ARI):** A different metric to compare the clusterings with the pre-defined cell-type labels. **Label silhouette width (ASW):** measures the within-cluster distance of cells compared to the distance to the closest neighboring cluster. A value close to 1 indicates a high separation between clusters. We used the predefined cell labels to define clusters for the label silhouette width calculation. **Graph cLISI:** measures the separation of the kNN graph. It evaluates the likelihood of observing the same cell-type label in the nearest neighbors, indicating good cell-type separation. **Isolated label metrics:** The isolated labels are defined as the cell types present in the fewest number of batches (Supplementary Table 1). Two metrics evaluate how well isolated labels separate from other cell types. The F1 score is the harmonic mean of precision and recall. The isolated label silhouette measures the ASW of the isolated label compared to all non-isolated labels. **Trajectory conservation:** Computes the correlation of inferred pseudotime ordering before and after integration.

Four metrics measure different levels of batch integration.

**Principal component regression:** measures the amount of variance of the principal components of the embedded space that can be explained by the batch variables before and after integration. **Graph connectivity:** measures whether the kNN graph of the embedding connects all cells that have the same cell type label. If there are strong batch effects, this will not be the case. **Graph iLISI:** measures the mixture of the kNN graph. It evaluates the likelihood of observing different batch labels in the nearest neighbors indicating a good batch separation. The **batch silhouette width** is a metric similar to the label silhouette width but applied to batch labels. To ensure that higher scores represent better mixing, the silhouette metric is subtracted from 1. The ASW is computed separately for each cell label to assess the mixing within cells of the same label. Finally, the individual ASW scores for each cell label are averaged to obtain an overall measure of batch mixing.

#### Enrichment analysis

Enrichment analysis was performed with respect to four sets of regulatory elements: distal enhancers, super-enhancers, highly expressed genes, and highly variable genes.

Annotations for distal enhancers in the hg38 genome assembly were downloaded from ENCODE Registry of CREs (v3, screen.encodeproject.org)^31^. They were then subset to distal cCREs with enhancer-like signatures (dELS) and CTCF-bound cCREs with enhancer-like signatures (CTCF-bound, dELS).

Super-enhancers were downloaded from SEdb 2.0 (http://www.licpathway.net/sedb/)^32^. Only bone marrow samples were included.

Highly expressed genes were computed using the preprocessed scRNA-seq data from the NeurIPS dataset. They were defined as the top 2000 genes ranked by mean expression across all cells.

Highly variable genes were computed with scanpy^33^ (v.1.9.2) using Seurat-based highly variable gene selection with default parameter settings.

We filtered annotations to overlap with at least one peak of the NeurIPS dataset. Region overlap was determined using the pyRanges package (v.0.0.124). Odds ratios and significance were computed using the Fisher exact test implemented in scipy (v.1.10.1) and corrected for multiple testing with Benjamini-Hochberg at an FDR of 0.05.

#### Correlation with gene expression analysis

We used the peak annotation of Cell Ranger ATAC to subset high-count peaks to promoter regions. Cell Ranger annotates a peak as a promoter if it overlaps with the promoter region (−1000 bp, +100 bp) of any transcription start site^17^. Then, we computed the Spearman correlation between a cell’s fragment count in the promoter peaks and the gene expression count using scipy, taking only cells with a fragment count > 1 into account. Since this correlation can be driven by cells with a high total fragment count, we restricted the computation to cells whose total fragment count was in the 0.25-0.75 quantile.

#### Normalized accessibility

We can use the learned latent space and generative model of Poisson VAE and Binary VAE to produce denoised and normalized estimates of accessibility, controlling for sequencing depth^25^. To this end, we defined as normalized accessibility of the model output using the median total fragment count across all cells. For cisTopic, we used the imputed and normalized accessibility scores.

We compared the normalized accessibility of the Poisson VAE model and the Binary VAE model by computing the cell type separation using the silhouette width (ASW) and area under the ROC curve.

## Data availability

Raw published data for the NeurIPS^24^, Satpathy^14^, the fly^15^, and the sci-ATAC-seq3^16^ datasets are available from the Gene Expression Omnibus under accession codes GSE194122, GSE129785, GSE163697, and GSE149683, respectively.

## Code availability

All models, code, and notebooks to reproduce our analysis and figures, as well as a tutorial notebook to use the Poisson VAE model, are available on https://github.com/theislab/scatac_poisson_reproducibility. The Poisson VAE model is available as an extension of the scvi-tools suite at https://github.com/lauradmartens/scvi-tools.

## Author contributions

L.D.M conducted the analysis and implemented the models. J.G. and F.J.T. conceived and supervised the project with the help of D.S.F. and V.A.Y. All authors wrote and contributed to the manuscript. The authors read and approved the final manuscript.

## Corresponding authors

Correspondence to Julien Gagneur or Fabian J. Theis.

## Acknowledgments

We would like to thank Ignacio L. Ibarra and Fabiola Curion for valuable feedback and Ignacio L. Ibarra, Alexander Karollus, Pedro Tomaz da Silva for feedback on the manuscript. L.D.M acknowledges support by the Helmholtz Association under the joint research school Munich School for Data Science - MUDS”, J.G by the Deutsche Forschungsgemeinschaft (SFB/Transregio TRR267). Fig. 1a is adapted from “ATAC Sequencing” by BioRender.com (2022). Fig. 2h is adapted from “Regulation of Transcription in Eukaryotic Cells” by BioRender.com (2022). Retrieved from https://app.biorender.com/biorender-templates.

