## Extended Data Figures for "Modeling fragment counts improves single-cell ATAC-seq analysis"

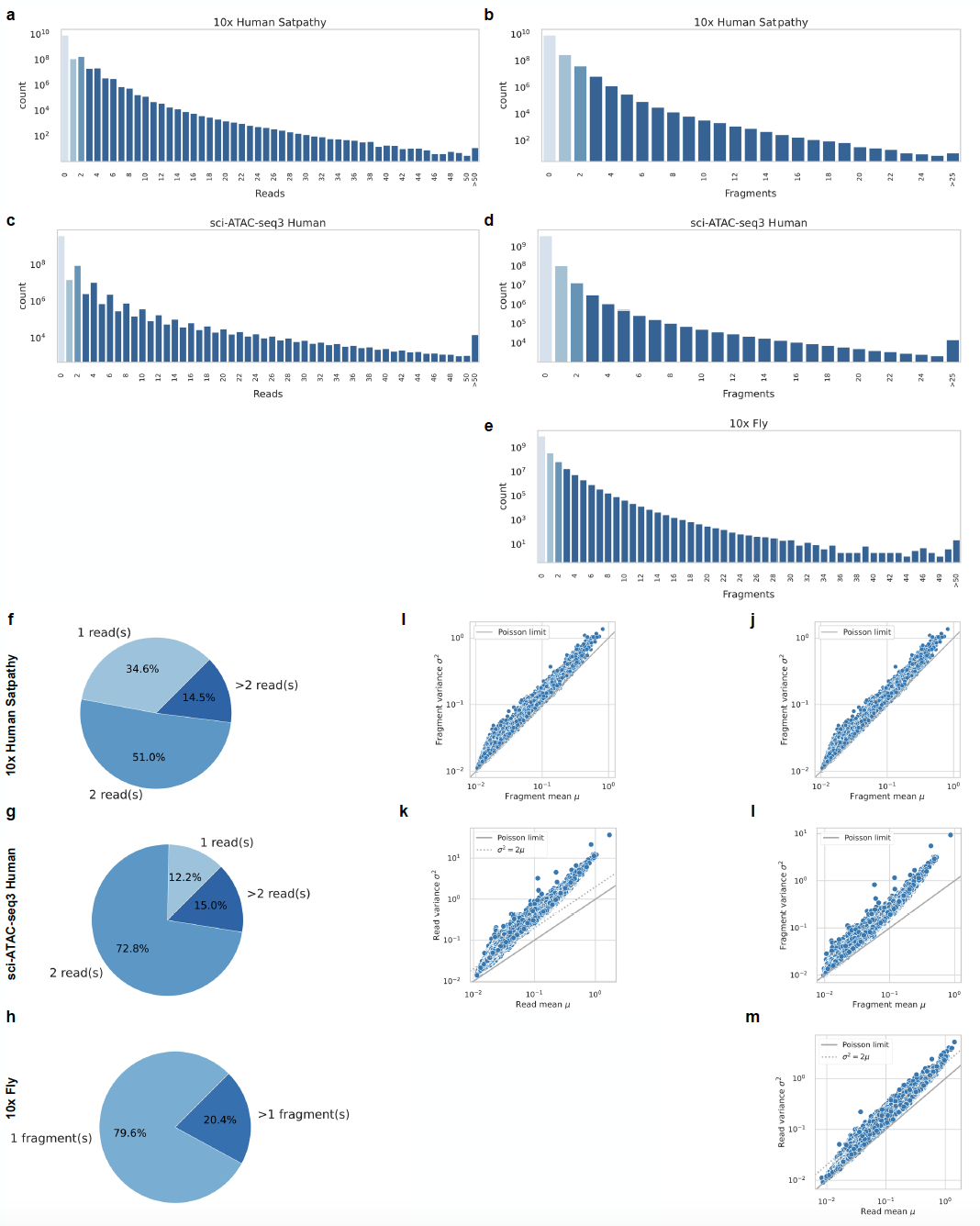

**Extended Data Figure 1**

**a, b)** Read count (a) and fragment count (b) distribution on the Satpathy dataset^1^. **c, d)** Read count (c) and fragment count (d) distribution of the sci-ATAC-seq3 dataset^2^. Plotted is a 10% random subset as the dataset consists of ~700K cells. **e)** Fragment count distribution on the fly dataset^3^. Cellranger ATAC read counts were unavailable for this dataset as we generated fragment counts directly with Signac. **f, g)** Pie chart showing the percentage of all non-zero peaks with 1, 2, or more than 2 reads for the Satpathy dataset (f), sciATAC-seq3 dataset (10% random subset) (g). **h)** Pie chart with the percentage of all non-zero peaks with 1 or more than 1 fragment for the fly dataset (read counts are not available for this dataset). **i, j)** Variance of read counts across cells against mean read counts for the Satpathy dataset (i) and sciATAC-seq3 dataset (j). Each dot represents one peak region. When fragment ends (reads) are counted, the variance of read counts is around twice the mean (grey dotted line), which is not consistent with a Poisson distribution (solid grey line). **k, l, m)** Same as (i, j), but for fragment counts.

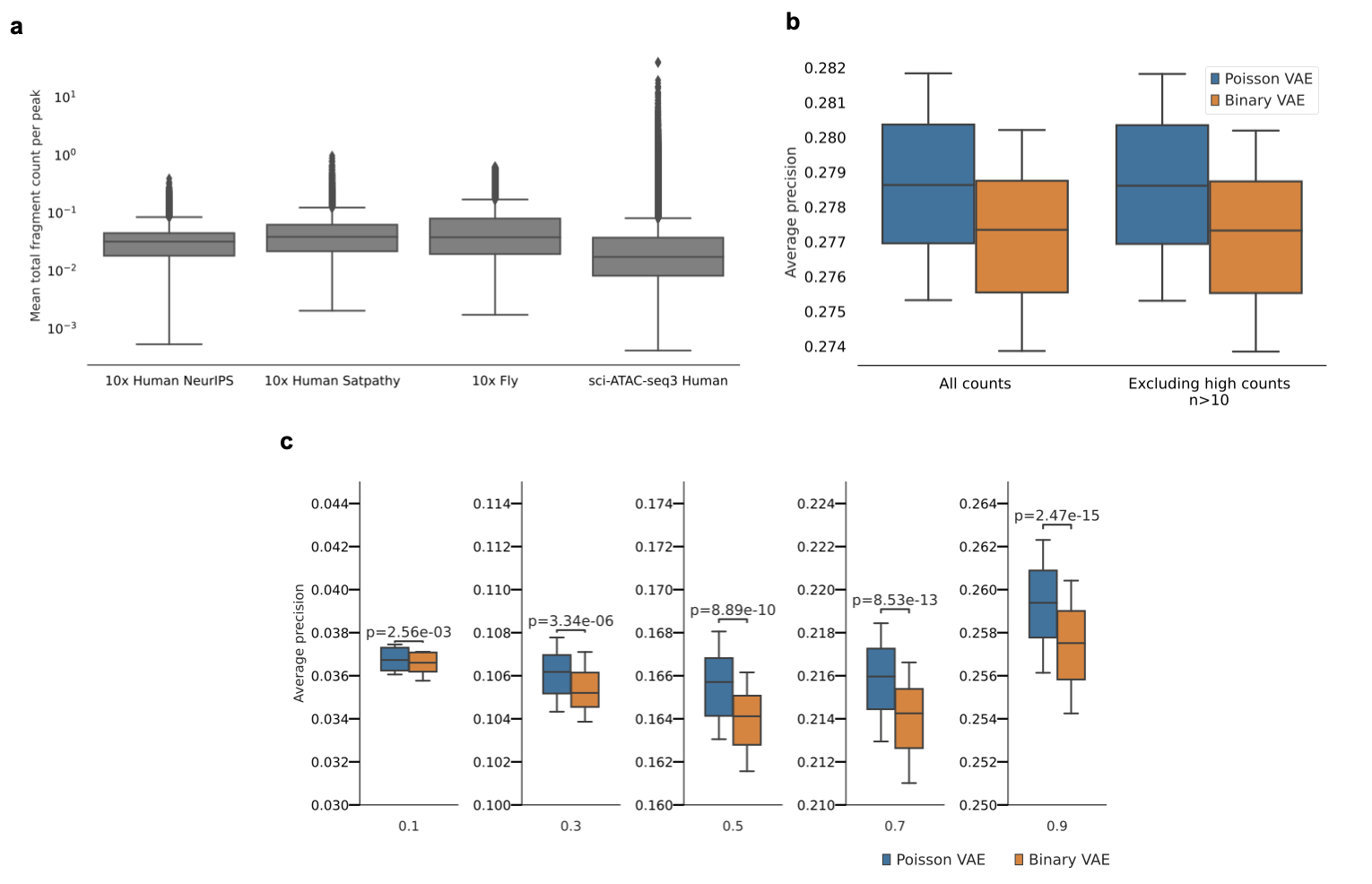

**Extended Data Figure 2**

**a)** Average fragment count distribution per peak for all four datasets. The sci-ATAC-seq3 dataset is 50% sparser than the 10x datasets. **b)** Average precision of the Poisson VAE and the Binary VAE model on the NeurIPS dataset for all cell-peaks and only the subset of cell-peaks that have less than 10 counts. **c)** Average precision for the Poisson VAE and Binary VAE model at different downsampling thresholds. *P* values were computed using the paired t-test. In boxplots, the central line denotes the median, boxes represent the interquartile range (IQR), and whiskers show the distribution except for outliers. Outliers are all points outside 1.5 times the IQR.

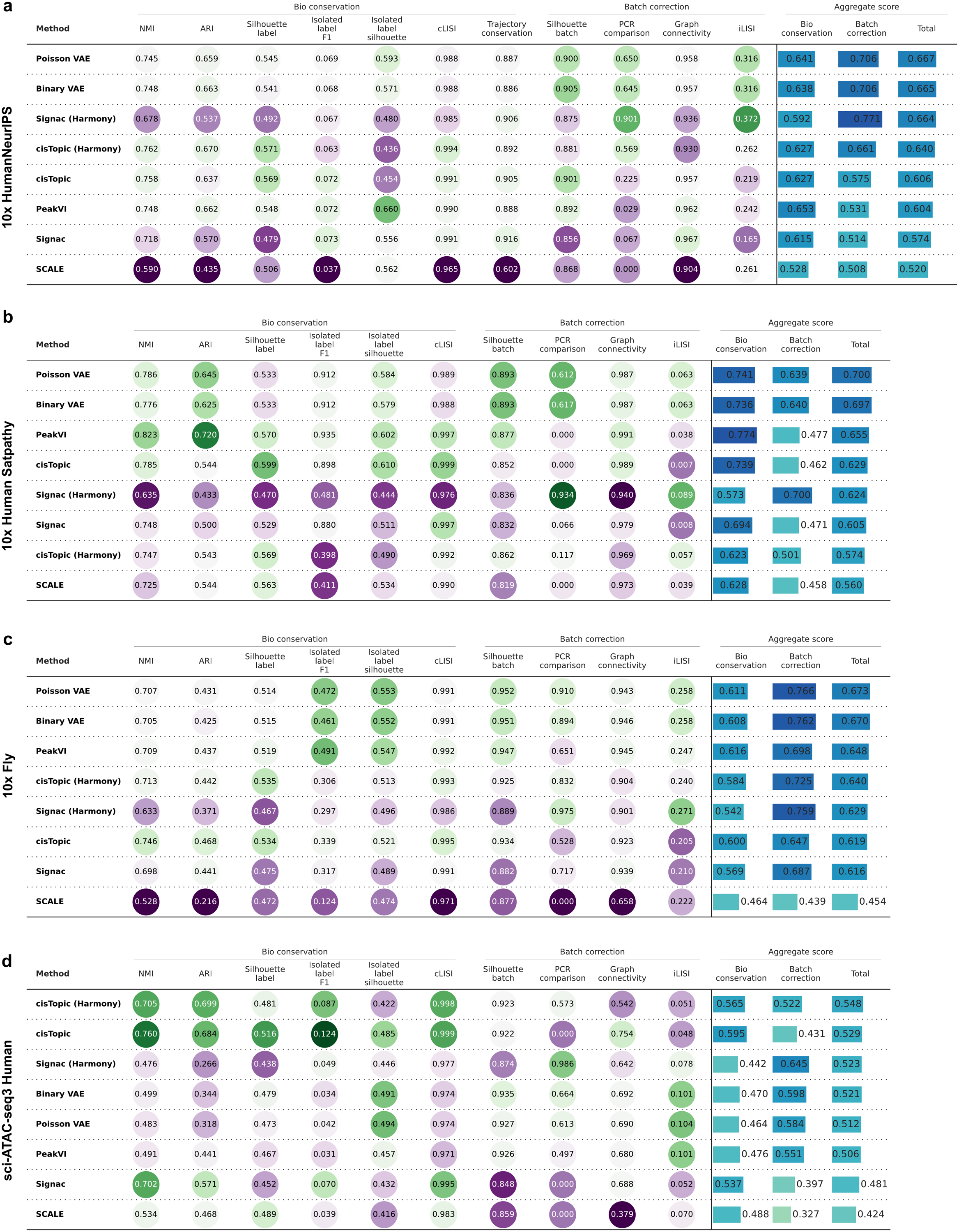

**Extended Data Figure 3**

Comparison of integration accuracy for Poisson VAE, Binary VAE, PeakVI^4^, Signac^5^ using LSI, CisTopic^6^ using LDA and SCALE^7^ on (**a**) the NeurIPS, (**b**) the Satpathy (**c**) the fly and (**d**) the sci-ATAC-seq3 datasets. For CisTopic and Signac, additional batch correction was performed using Harmony. Metrics are categorized into batch correction (blue) and bio-conservation (pink) categories. Reported is the mean over 10 cross-validation runs. Overall scores were computed using a 40:60-weighted mean of batch correction and bio-conservation scores.

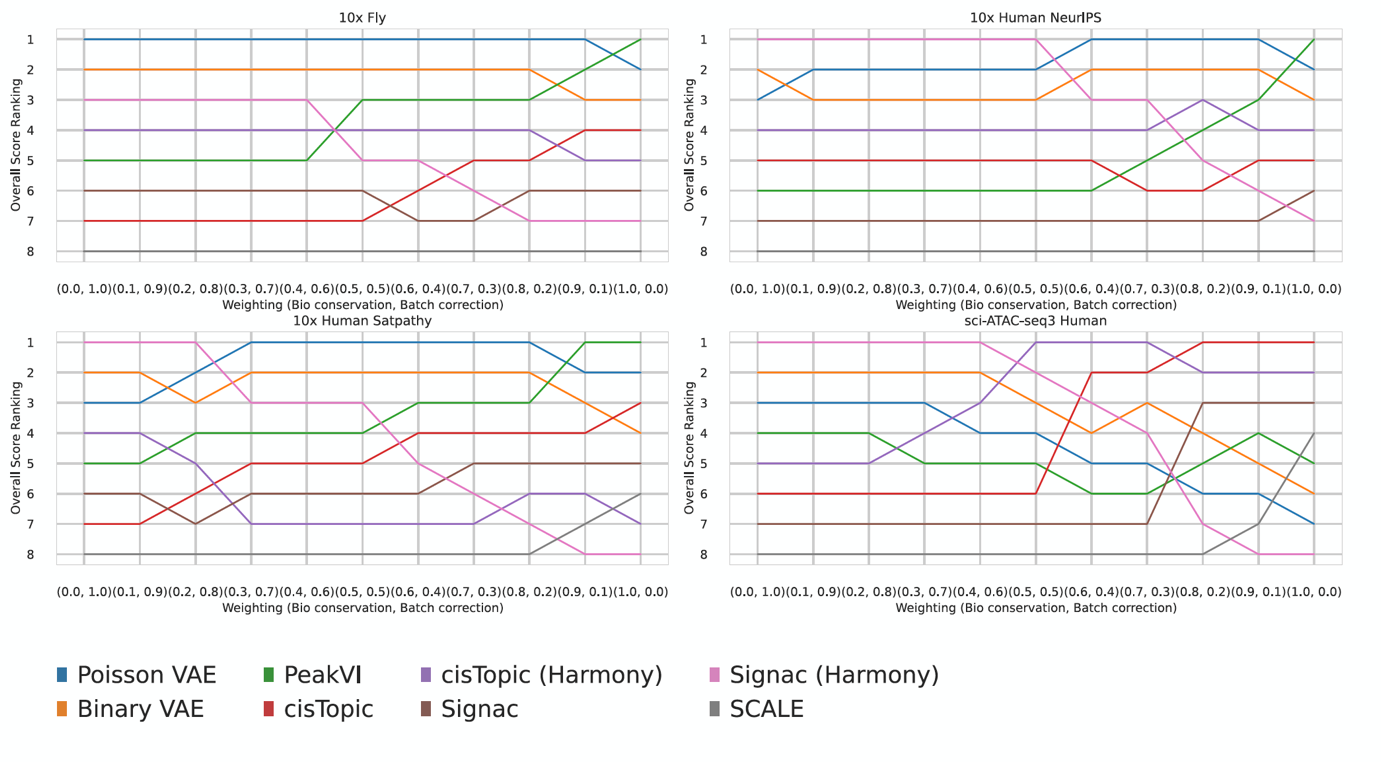
**Extended Data Figure 4**

Overall score computed from different weightings of bio conservation and batch correction.

**Extended Data Figure 5: NeurIPS dataset**

UMAP of the integrated latent space of the NeurIPS dataset using the Poisson VAE, Binary VAE, PeakVI, Signac using LSI, CisTopic using LDA, and SCALE model. Cells are colored by cell type (top row) and batch (bottom row). For CisTopic and Signac, additional batch correction was performed using Harmony.

**Extended Data Figure 6: Satpathy datset**

UMAP of the integrated latent space of the Satpathy dataset using the Poisson VAE, Binary VAE, PeakVI, Signac using LSI, CisTopic using LDA, and SCALE model. Cells are colored by cell type (top row) and batch (bottom row). For CisTopic and Signac, additional batch correction was performed using Harmony.

**Extended Data Figure 7: Fly dataset**

UMAP of the integrated latent space of the fly dataset using the Poisson VAE, Binary VAE, PeakVI, Signac using LSI, CisTopic using LDA, and SCALE model. Cells are colored by cell type (top row) and batch (bottom row). For CisTopic and Signac, additional batch correction was performed using Harmony.

**Extended Data Figure 8: sci-ATAC-seq3 dataset**

UMAP of the integrated latent space of the sciATAC-seq3 dataset using the Poisson VAE, Binary VAE, PeakVI, Signac using LSI, CisTopic using LDA, and SCALE model. Cells are colored by cell type (top row) and batch (bottom row). For CisTopic and Signac, additional batch correction was performed using Harmony.

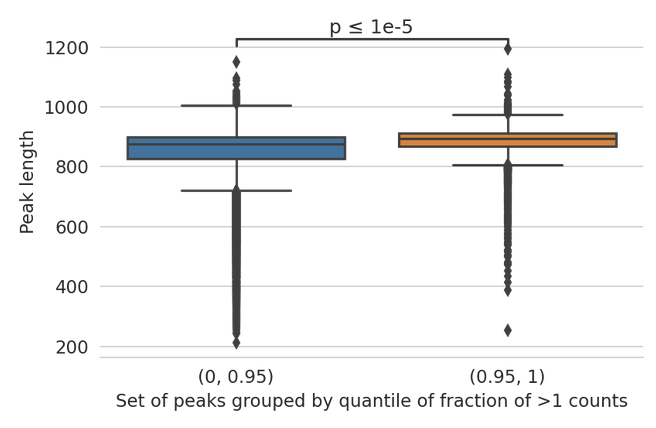

**Extended Data Figure 9**

Peak distribution length for peaks in the top 0.05 quantile and bottom 0-0.95 quantile according to the fraction of counts above the binarization threshold. High-count peaks are significantly longer. The central line denotes the median, boxes represent the interquartile range (IQR), and whiskers show the distribution except for outliers. Outliers are all points outside 1.5 times the IQR. The *P* value was computed using a Wilcoxon test.

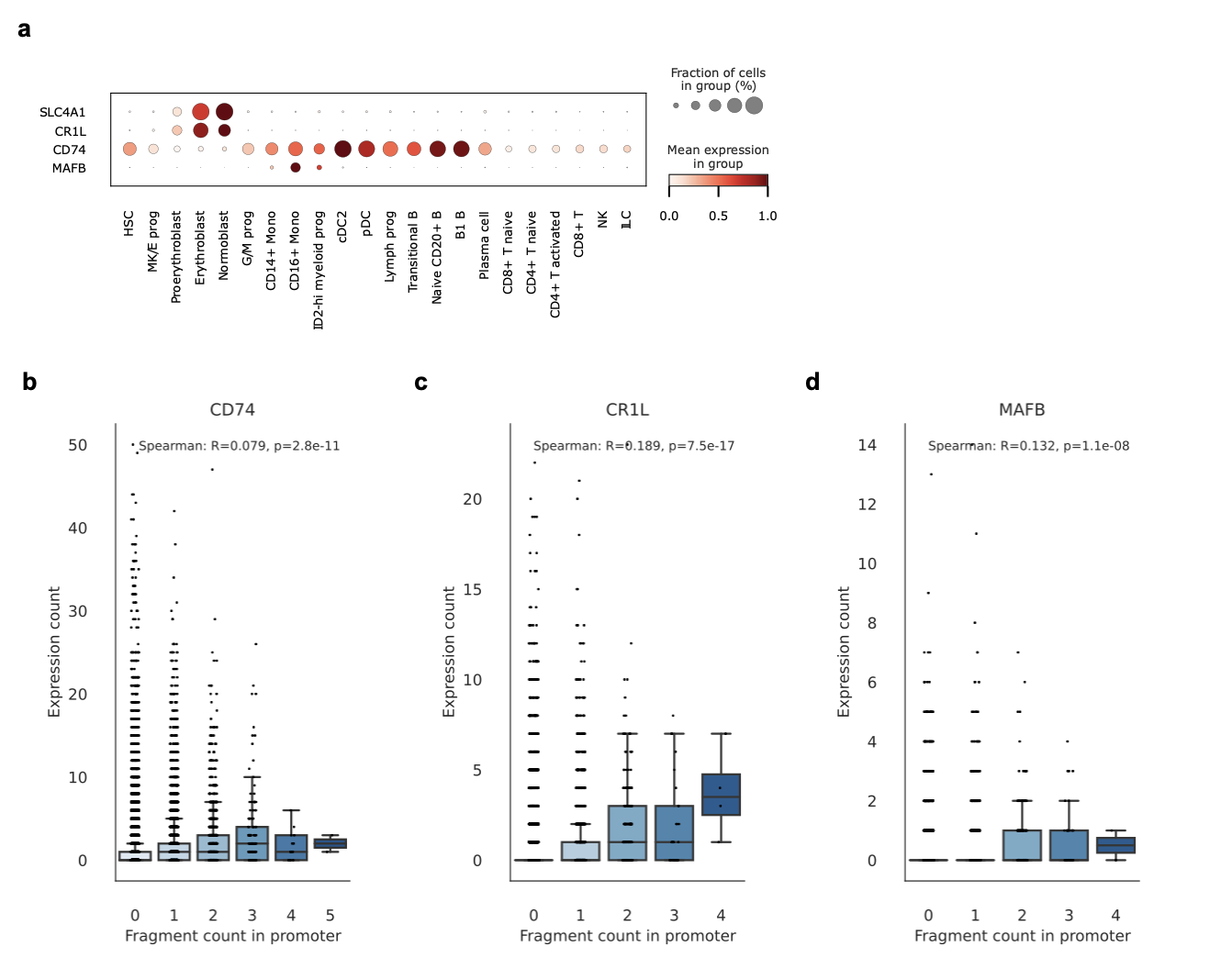

**Extended Data Figure 10**

**a)** Expression of genes (rows) associated with each cell type (columns). *SLC4A1* and *CR1L* are involved in the red blood cell lineage^9,10^ (Proerythroblast, Erythroblast, Normoblast). *CD74* is expressed in antigen-presenting cells and is known to regulate mature B-cell survival^11^. *MAFB* is a transcription factor that represses erythrocyte programs in myeloid cells^12^. Correlation of gene expression and fragment counts in the promoter of the (**b**) *CD74* gene, (**c**) *CR1L* gene, and (**d**) *MAFB* gene. The Spearman correlation was computed on fragment counts greater than 0. We restricted the plot to cells of similar total fragment count (0.25-0.75 quantile) to avoid capturing effects driven by total fragment count. We see a quantitative signal in promoter accessibility that would be lost by binarization.

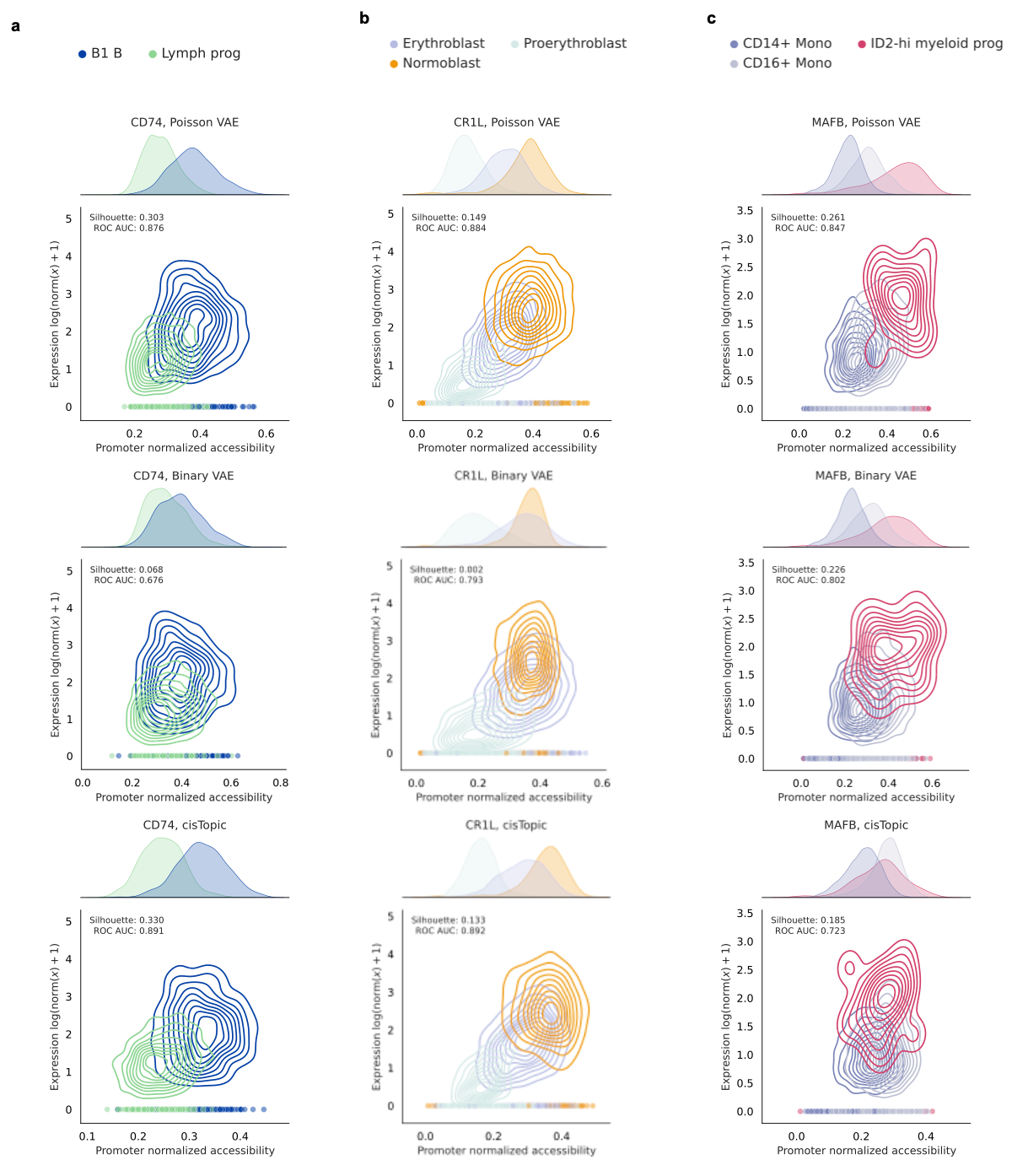
**Extended Data Figure 11**

**a, b, c)** Log-normalized gene expression against normalized accessibility for the Poisson VAE (top row) , Binary VAE model (middle row) and cisTopic model (bottom row) for the (**a**) *CD74* gene, (**b**) *CR1L* gene, and (**c**) *MAFB* gene. Cell type separation is measured with the silhouette width and area under the ROC curve and is better with the Poisson VAE model for CR1L and MAFB and second for CD74.
